# Minimized Sample Consumption for Time-Resolved Serial Crystallography Applied to the Redox Cycle of Human NQO1

**DOI:** 10.1101/2024.04.29.591466

**Authors:** Diandra Doppler, Alice Grieco, Domin Koh, Abhik Manna, Adil Ansari, Roberto Alvarez, Konstantinos Karpos, Hung Le, Mukul Sonker, Gihan K. Ketawala, Samira Mahmud, Isabel Quereda Moraleda, Angel L. Pey, Romain Letrun, Katerina Dörner, Jayanath C. P. Koliyadu, Raphael de Wijn, Johan Bielecki, Huijong Han, Chan Kim, Faisal H. M. Koua, Adam Round, Abhisakh Sarma, Tokushi Sato, Christina Schmidt, Mohammad Vakili, Dmitrii Zabelskii, Richard Bean, Adrian P. Mancuso, Joachim Schulz, Raimund Fromme, Milagros Medina, Thomas D. Grant, Petra Fromme, Richard A. Kirian, Sabine Botha, Jose Manuel Martin-Garcia, Alexandra Ros

## Abstract

Sample consumption for serial femtosecond crystallography (SFX) with X-ray free electron lasers (XFELs) remains a major limitation preventing broader use of this powerful technology in macromolecular crystallography. This drawback is exacerbated in the case of time-resolved (TR)-SFX experiments, where the amount of sample required per reaction time point is multiplied by the number of time points investigated. Thus, in order to reduce the limitation of sample consumption, here we demonstrate the implementation of segmented droplet generation in conjunction with a mix-and-inject approach for TR studies on NAD(P)H:quinone oxidoreductase 1 (NQO1). We present the design and application of mix-and-inject segmented droplet injectors for the Single Particles, Clusters, and Biomolecules & Serial Femtosecond Crystallography (SPB/SFX) instrument at the European XFEL (EuXFEL) with a synchronized droplet injection approach that allows liquid phase protein crystal injection. We carried out TR-crystallography experiments with this approach for a 305 ms and a 1190 ms time point in the reaction of NQO1 with its coenzyme NADH. With this successful TR-SFX approach, up to 97% of the sample has been conserved compared to continuous crystal suspension injection with a gas dynamic virtual nozzle. Furthermore, the obtained structural information for the reaction of NQO1 with NADH is an important part of the future elucidation of the reaction mechanism of this crucial therapeutic enzyme.

## Introduction

Serial Crystallography both at X-ray free electron lasers (XFELs) and synchrotrons has developed into a robust tool for protein structural analysis. It surpasses limitations in traditional goniometer-based crystallography approaches requiring large crystals where X-ray damage can be prohibitive and cryogenic approaches remain imperative. Similarly, crystal defects become problematic in larger crystals required for traditional crystallography approaches, whereas in SFX with XFELs, such effects remain suppressed as crystals in the order of 1-20 µm are typically employed.

Furthermore, serial crystallography has allowed dynamic studies on proteins both for light-induced reactions as well as those induced through a substrate or binding partner.^1^ To facilitate the latter approach termed mix-and-inject serial crystallography (MISC), the small crystals need to be mixed with a substrate or binding partner on time scales of milliseconds to initiate the reaction and then be probed by the X-ray beam at a certain time delay representing the time along the reaction coordinate.^2^ Time-resolved experiments with MISC have been carried out to study enzymatic reactions occurring from milliseconds up to seconds^3, 4^ with unprecedented results. Some examples, among many, of this dynamic SFX technique include the reaction of the β-lactamase C (BlaC) from *Mycobacterium tuberculosis* with the antibiotic ceftriaxone^5, 6^ and the inhibitor sulbactam^7^, the reaction of a riboswitch RNA with the substrate adenine^8^, the cytochrome C oxidase with O_2_^9^ and the isocyanide hydratase (ICH) enzyme, which catalyzes the hydration of isocyanides.^10^

Despite its success, a bottleneck in MISC that remains is the large amount of sample required to probe just enough snapshots of each reaction time point to obtain suitable information about the reaction mechanism. For every time point measured in a TR experiment with MISC, the amount of consumed sample increases to the extent necessary to obtain a complete data set, which typically amounts to a total of several hundreds of milligrams of protein. These quantities are often prohibiting TR-SFX with the MISC approach, as proteins may not be available in such large abundance. Therefore, past efforts in the improvement of serial crystallography techniques have targeted sample injection and delivery approaches that allow a significant reduction in the amount of sample needed for a full data set.

To this end, several approaches that allow the reduction of sample required for an SFX experiment have been described in the past years, which have been reviewed by Cheng *et. al*.^11^ One of these approaches is the fixed-target approach in which a crystal slurry loaded onto a thin support to minimize background, are scanned in front of the X-ray beam.^12^ They have been successfully employed, and typically require only a few microliters (< 1 mg of protein) of a crystal slurry to fill a chip from which a complete data set can be obtained. However, fixed-target devices may suffer from poor vacuum compatibility inducing dehydration, specifically when crystals are loaded between very thin support layers to reduce background or on non-enclosed devices to facilitate loading through wicking.^11^ Additionally, fixed target devices are limited in the speed at which the chip can raster in front of the X-ray source, therefore they are compatible with lower repetition rates. More importantly, since crystals are typically placed in an enclosed environment, fixed-target devices are impractical for TR-SFX experiments that require substrate mixing.

To sustain the advantages of established jet-based injection techniques, two main approaches that facilitate the reduction of sample consumption have been developed. The first approach are the viscous injectors which allow the injection of crystal samples in a highly-viscous stream, extruding samples at flow rates ranging from just a few nL/min up to only a few µL/min. Originally developed for SFX experiments with membrane protein crystals grown in lipidic cubic phase, these injectors have also been successfully tested to inject soluble protein crystals with a variety of viscous media^13–18^ both at XFELs and synchrotron radiation sources. An advantage of the viscous jets is that they can be coupled with a laser source to perform TR experiments for light-sensitive proteins, however, they are also limited in the speed of extrusion and are therefore only compatible with lower repetition rates.^19^ The second jet-based approach relates to the formation of droplets, ejecting them at a desired frequency to intersect with the X-ray beam. Free-standing droplets can be generated from nozzles with piezoelectric or acoustic actuation. The droplets generated with these approaches are in the range of pL-nL, significantly reducing sample requirements. However, injection into a vacuum remains challenging, and clogging effects may hamper crystallography for samples injected in droplets.

Another special class of droplet generators divides sample-laden droplets within an immiscible oil stream. These techniques not only facilitate *in situ* droplet crystallization before X-ray diffraction but also enable crystal-containing droplets to mix with a substrate through droplet coalescence post-crystal formation, thereby enabling time-resolved diffraction experiments. ^20^ Moreover, they are applicable for certain light-induced time-resolved studies and experiments requiring crystal injection into a low background environment (vacuum). They are particularly useful when crystals cannot grow to the appropriate size and quality required in viscous injection media or when other experimental parameters hinder the use of fixed-target approaches. Additionally, these techniques are particularly suited for experiments where liquid injection with a gas dynamic virtual nozzle (GDVN) is required. An intrinsic advantage of segmented flow droplet injection is that principles of continuous injection with a GDVN can directly be applied, such as established crystallization parameters (e.g. maintaining growth media and established crystal size), but also the characteristics of the created jet (such as jet velocity and jet thickness). Additionally, with small modifications, these devices can generate droplets that are compatible with both high and low-repetition rate XFELs. GDVN injection is a robust and still developing technique that has contributed to 30% of the structures added to the Protein Data Bank (PDB) using reported serial femtosecond crystallography experiments, thus investigating further development is imperative for propelling forward time-resolved structural biology, opening doors to unprecedented insights into dynamic molecular processes.^21^

SFX with the segmented droplet generation approach has indeed been demonstrated for several proteins so far, both at the Macromolecular Crystallography Instrument (MFX) at the Linac Coherent Light Source (LCLS) at SLAC National Accelerator Laboratory, and the European XFEL (EuXFEL), where sample savings of 75% and 60% have been demonstrated, respectively.^22–24^ However, a time-resolved crystallography experiment with segmented droplet injection has not been demonstrated to date. We estimate that the segmented droplet approach may reduce sample consumption by a factor of 100 at the EuXFEL when droplets of the size spanning an entire pulse train are created at 10 Hz. Segmented droplet injection would thus be ideally suited for time-resolved experiments based on the mix-and-inject principle at the EuXFEL.^25, 26^ Thus, it is our goal to establish a droplet injector with minimal sample consumption that would be available for routine user operation in TR-SFX experiments.

As a proof-of-concept experiment and to demonstrate that our novel device is suitable for TR-SFX experiments at the EuXFEL with reduced sample consumption, we have employed the human NAD(P)H: quinone oxidoreductase 1 (NQO1) and its coenzyme NADH. In a recent study, we have determined the first room-temperature structure of NQO1 in complex with NADH using serial synchrotron crystallography in combination with molecular dynamics simulations.^27^ Although relevant information was discovered on NQO1 dynamics and mechanism, the solved structure was a static picture of the reaction with NADH. Thus, to fully unravel the NQO1 function, exploration beyond static structures is required. To this end, in the work presented here, we have conducted the first TR-SFX experiments on the NQO1 enzyme with its coenzyme NADH at the EuXFEL. Two time points were investigated, 305 and 1190 ms, using a novel mix-and-inject segmented droplet injector. The previously established segmented droplet injection principle was extended to include a mixer element just upstream of the droplet generation region in one completely 3D-printed device. We demonstrated the integration into the SPB/SFX experimental chamber at the EuXFEL and applied the mixer/droplet injector to TR-SFX with NQO1 successfully. We focus on the conditions of optimized synchronization of the droplets with the XFEL pulse trains as well as the characterization of the reaction time points, and mixing times achieved in this approach. Our efforts are motivated by providing a sample conservation approach for SFX at the EuXFEL to take full advantage of the capabilities of this XFEL operating with MHz pulse trains repeating at 10 Hz frequency.

## Materials and Methods

### Materials

Perfluorodecalin (PFD) and 1H,1H,2H,2H-perfluoro-1-octanol (perfluorooctanol, PFO) were purchased from Sigma-Aldrich (St. Louis, USA). SU-8 developer was obtained from Microchem (Round Rock, USA). The photoresist IP-S was purchased from Nanoscribe GmbH (Eggenstein-Leopoldshafen, Germany). Deionized water (18 MΩ) was supplied from an LA755 Elga purification system (Elga Lab water, High Wycombe, USA) and isopropyl alcohol (IPA) and ethanol were obtained from VWR International (Radnor, USA) or Decon Labs (King of Prussia, USA). Fused silica capillaries (360 µm outer diameter, 100 µm inner diameter) were purchased from Molex (Lisle, USA). Hardman extra-fast setting epoxy glue was purchased from All-Spec (Houston, USA).

*E. coli* BL21 (DE3) competent cells were purchased from Agilent technologies (USA). Yeast extract and tryptone were purchased from Condalab (Madrid, Spain). EDTA-free Protease Inhibitor Cocktail, Isopropyl β-D-1-thiogalactopyranoside (IPTG), ampicillin, sodium phosphate, sodium chloride, imidazole, flavin adenine dinucleotide (FAD), sodium acetate, K-HEPES, and Tris-HCl were purchased from Merck (Madrid, Spain). Polyethylene glycol (PEG) 3350 was purchased from Hampton Research (USA), and nicotinamide adenine dinucleotide (NADH) was purchased from Merck (Madrid, Spain).

### Mixer Device Design and Fabrication

Droplet generation devices that facilitate mixing with a substrate for time-resolved studies were employed during the experiments. These devices were comprised of three components, as detailed in **Figure 1a**. The first component integrated a droplet generator with a Y-shaped Mixer and also featured integrated electrodes as described previously.^22, 23^ The second component consisted of an optical fiber holder, and the third was a 3D printed GDVN. All components were connected with joining fused-silica capillaries and glued together using epoxy glue.

**Figure 1:**
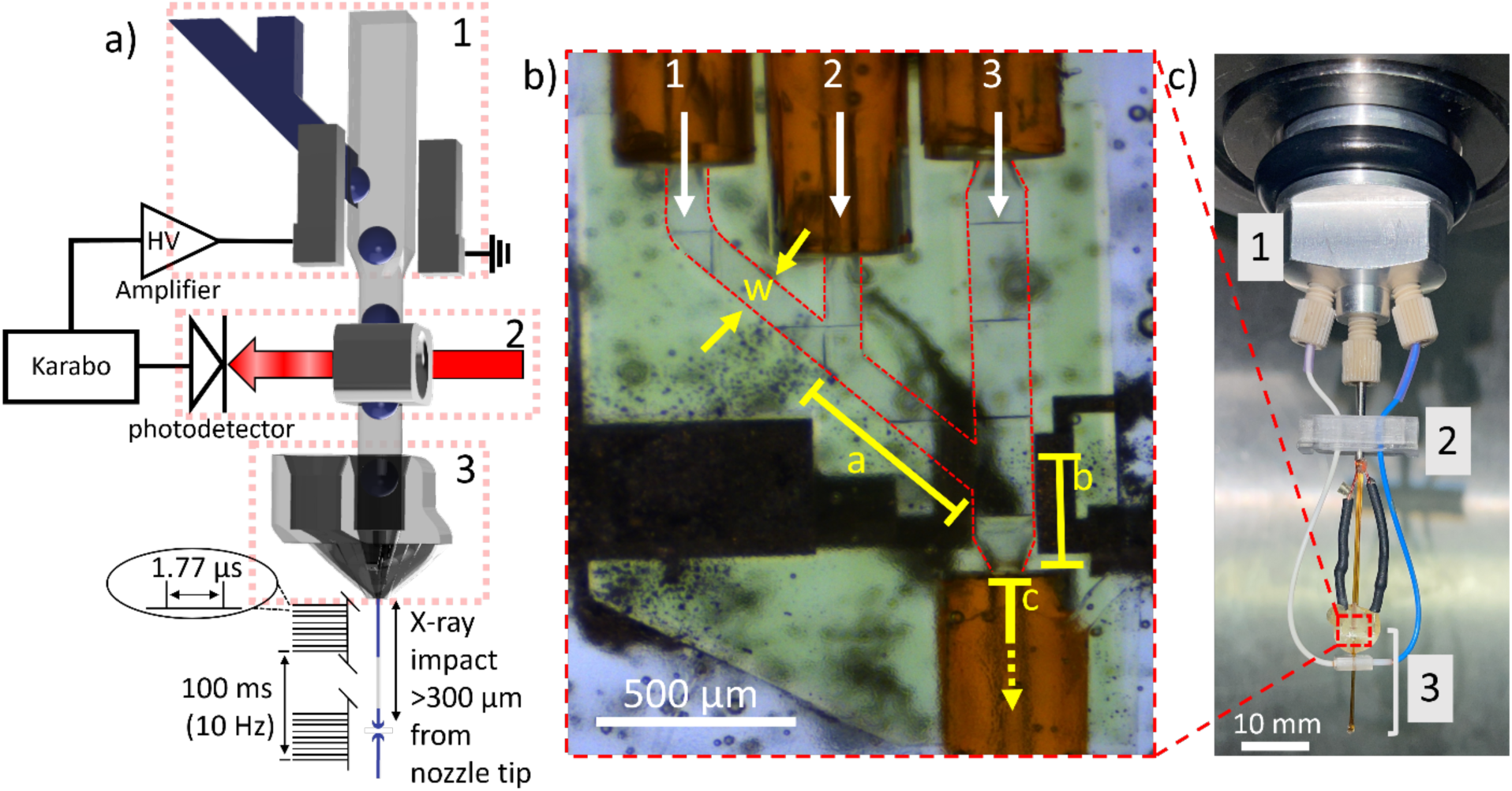
**(a)** Schematic representation of the droplet generator setup with control hardware and X-ray interaction region in the SPB/SFX instrument at the EuXFEL highlighting the following components: 1) Y-Mixer droplet generator with integrated electrodes, 2) optical fiber droplet detector, and 3) 3D printed gas dynamic virtual nozzle. **(b)** Microscopy image of fully assembled DG250-Y-Mixer, illustrating the delivery capillaries of substrate (1), protein crystals (2), oil (3), and the exit point for mixed droplets (4). Critical dimensions are highlighted in yellow: w = 100 µm (width of the aqueous channel), a = 528 µm (length of channel section A within the mixer where substrate and crystal streams first mix), b = 285 µm (length of the channel section B where aqueous streams form a droplet segmented by the oil phase), and c = 15.4 mm (length of the remaining device, capillary, and region in the nozzle to the orifice of the GDVN in section C (not shown in full in the image)). **(c)** Injector mounted on the EuXFEL nozzle rod, featuring a custom adapter (1) and fiber bracket (2) securing the device and fibers (3) during insertion and experimentation.

The components of the droplet generator and integrated mixers were designed and fabricated following established procedures.^28^ Briefly, the devices were created using Fusion 360 (AutoDesk, San Francisco, USA) and subsequently 3D printed with a Photonic Professional GT 3D printer (Nanoscribe GmbH, Eggenstein-Leopoldshafen, Germany), employing IP-S photoresist. After printing, the devices underwent development in SU-8 developer followed by rinsing in IPA. Similar protocols were also employed for the GDVNs.

The droplet generator with an integrated mixer, named the DG-Y-Mixer, was comprised of three rectangular cross-section channels that converged in a series of Y-junctions, shown as a CAD rendering and a microscopy image, respectively, in **Figure 1a-b**. The continuous channel delivered the oil phase that segmented the droplets, the center channel delivered the protein crystals suspended in their mother liquor, and the channel farthest from the oil channel delivered the substrate dissolved in the same mother liquor. The oil phase was a 10:1 v/v mixture of PFD to PFO.

Two different versions of the DG-Y-Mixer were used during the experiments. The DG250-Y-Mixer, depicted in **Figure 1a-c**, contained aqueous channels that were 100 µm x 100 µm in cross-section that joined an oil channel with a cross-section of 150 µm x 150 µm. The second version, the DG300-Y-Mixer, depicted in **Figure 2c-d**, contained channels of all the same cross-section of 150 µm x 150 µm.

**Figure 2:**
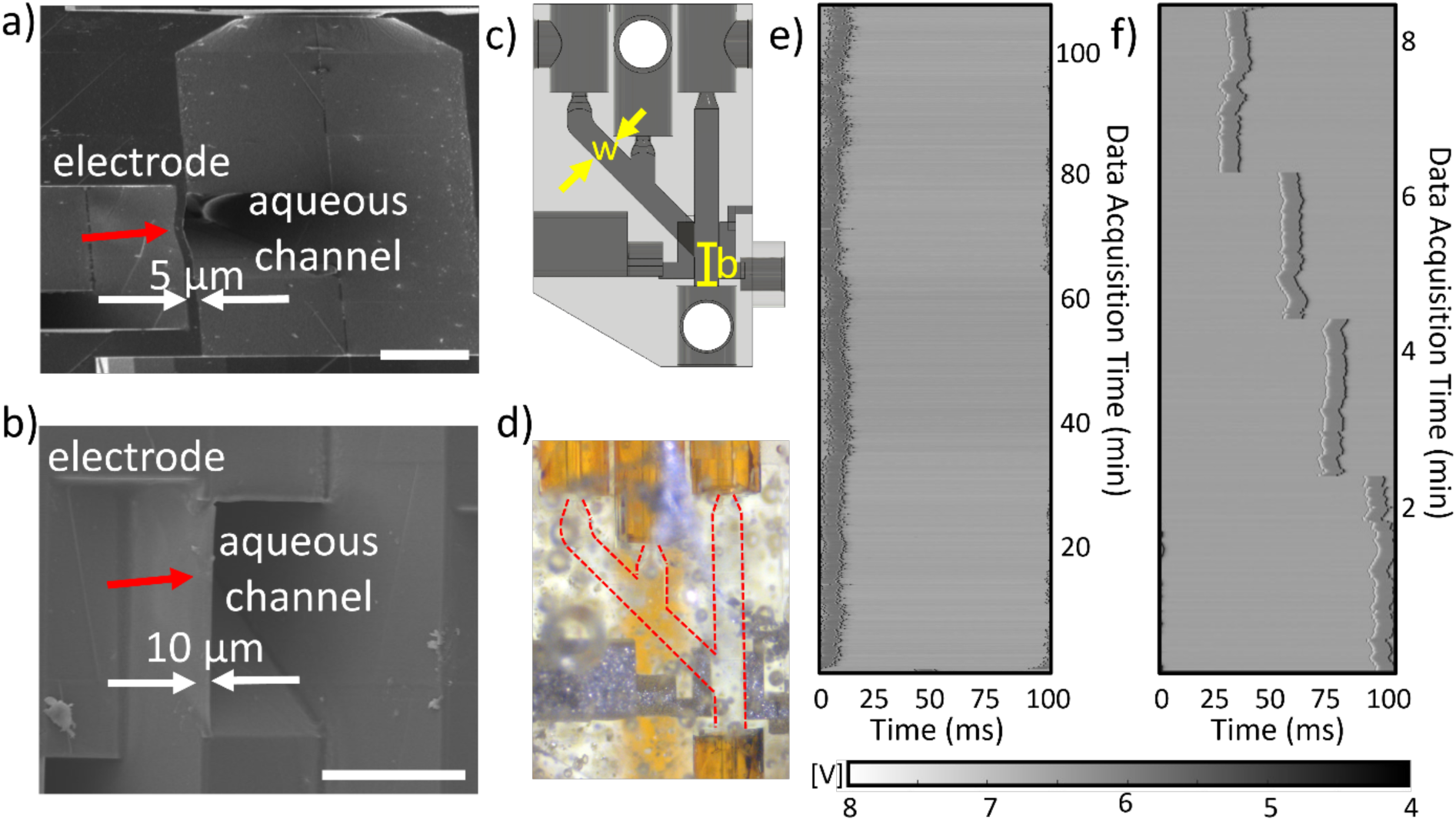
a) SEM image of the internal structure of the DG300 where a deformation in the 5 µm wall is indicated in red between the aqueous and the electrode channels. b) SEM image of the internal structure of the updated DG300 where the same channel, now 10 µm thick, is indicated in red, but not deformed. Aspect ratios in each SEM image differ and the scale bar represents 200 µm. c) CAD image and d) microscopy image of the DG300-Y-Mixer implemented at the EuXFEL. e) Resulting waterfall plot in a droplet generation device without a mixer in DG300 geometry, with *Q_O_* = 18.2 µL/min, *Q_B_* = 1.5 µL/min, trigger amplitude of 100 V and duration of 10 ms and f) waterfall plot from a DG300 at *Q_O_* = 18.5 µL/min, *Q_B_* = 1 µL/min, trigger amplitude of 200 V and duration of 10 ms with various triggering phases after the improvements in geometry and surface treatment. To establish the waterfall plot in e) and f), the droplet detector signal is divided into 100 ms sections according to the XFEL-reference at 10 Hz, and each trace is aligned in a vertical stack with the color representing the amplitude of the signal.

### SEM Imaging

Scanning electron microscopy (SEM) imaging was carried out at the Eyring Materials Center at Arizona State University, using the Zeiss Auriga FIB/SEM (Germany). Devices were printed with an open structure in the region of interest and then sputter-coated with gold. The SEM instrument was operated at 5.0 kV and the sample was tilted to inspect the wall between electrode and fluid channels.

### Fluidic Operation and SPB/SFX Configuration

Oil and crystal samples were driven from custom stainless-steel reservoirs by HPLC pumps (LC20AD, Shimadzu Co., Kyoto, Japan) or syringe pumps (neMESYS 1000, Cetoni, Korbussen, Germany). Bronkhorst liquid-flow sensors (mini CORI-FLOW™ ML120V00, Bronkhorst, Bethlehem, USA, and LIQUI-FLOW™ Mini, Bronkhorst, Bethlehem, USA) monitored flow rates in liquid lines before the reservoirs. PEEK tubing (Zeus, Orangeburg, USA, 250 μm ID, and 1/16-in OD), along with fittings and ferrules from IDEX Health & Science LLC (Oak Harbor, USA), connected the sensors, reservoirs, and tubing. Fused silica capillaries connected the reservoirs and the droplet generators.

For beam times P4502 and P3083 at the SPB/SFX instrument of the EuXFEL, devices and fibers were mounted on a custom-made adapter at the end of a nozzle rod and lowered into the vacuum chamber through the instrument’s load-lock system.^29^ Capillaries and insulated wires were fed through a steel tube affixed to the center of the adapter, while fibers were threaded through two additional side ports, as depicted in **Figure 1c**. The stainless-steel reservoirs holding the protein crystal suspension were mounted on a rotating shaker device (PTR-360, Radnor, VWR). Helium pressure for the GDVN was regulated by a GP1 electronic pressure regulator (Equilibar, Fletcher, USA) and monitored by a Bronkhorst mass flow sensor (EL-FLOW Select F-111B, Bronkhorst, Bethlehem, USA).

### Feedback Mechanism

During experiments conducted at the EuXFEL, the droplet monitoring and triggering feedback mechanism were fully integrated into Karabo, the in-house control system.^30^ This triggering scheme resembles our previous approach with the Raspberry Pi;^22, 23^ however, with all controls integrated, the droplet trace, recorded by a SIS8300 digitizer (Struck Innovative Systeme GmbH, Hamburg, Germany), and additional diagnostic information were recorded and saved within the EuXFEL data stream for post-experimental processing.^22, 23^

### Numerical Modeling

Numerical modeling was used to assess the spread of the reaction time point with COMSOL Multiphysics Ver. 6.2 (COMSOL, Burlington MA, USA). A 2D model of the of the aqueous channel was employed to simulate the diffusion for the substrate molecule in the convective flow in *section A.* To assess the substrate concentration in the droplet forming at the Y-intersection in *section B*, the geometry corresponded to the aqueous sample channel for either the DG250-Y-Mixer or DG300-Y-Mixer and the oil channel joined at a 45° angle designed in AutoCAD (AutoDesk, San Francisco, CA, USA).

The 2D simulation of *section A* utilized a Laminar Flow module (for pressure-driven flow) to establish the flow profile and the Transport of Diluted Species module for modeling convection and diffusion. The geometry used for modeling droplet formation contained both *section A* and *section B*, with the dimensions outlined in **Table SI-3**. The Laminar Flow module, employing the Level Set module for two-phase flow for droplet generation, and the Transport of Diluted Species module to model diffusion-convection mixing were employed. A mesh was created with the “extremely fine” setting applied over the full length of the channel and the duration of the study was 1.35 s with 1 ms step size for the DG300-Y and 0.78 s with 0.5 ms step size for the DG250-Y geometry, respectively. The model was built as previously reported ^31^ but modified for the presented channel geometries and experimentally employed liquid phase parameters. The details of boundary conditions, relevant equations, and parameters are detailed in **Table SI-3**.

### Protein and Crystal Sample Preparation

Protein expression and purification of human NQO1 were carried out as previously described.^22^ For beam time P3083, microcrystals of the free NQO1 were obtained on-site in the XBI labs of the EuXFEL using the batch with agitation method as follows: in a 3 mL glass vial, 100 μL of the protein solution at 18 mg/mL were slowly added dropwise to 300 μL of the precipitant solution (0.1 M Tris pH 8.5, 0.2 M sodium acetate, 25 % polyethylene glycol (PEG) 3350) while stirring at 200 rpm. Upon addition of the protein, the solution turned turbid immediately and needle-shaped crystals of an average size of 40 µm in their longest dimension grew at room temperature in about 1 h. Before loading the sample into the reservoir, the microcrystal samples were filtered through an inline filter (20 µm pore size) and then concentrated four times by centrifugation at 3,000 rpm for 2 min to settle crystals. In the case of beam time P4502, a slightly different procedure was applied to produce the crystal sample. Smaller microcrystals of 10-20 µm in their longest dimension were grown on-site by adding a protein solution at a higher concentration (26.5 mg/mL). Before loading the sample into the reservoir, the microcrystal samples were filtered through an in-line filter with a mesh cut-off size of 26 µm x 26 µm and further concentrated four times by letting them settle for about 4 h and subsequently removing the supernatant. The final crystal concentration for the experiments was 2 x 10^7^ crystals/mL in P3083 and 2 x 10^9^ crystals/mL in P4502. For the mixing experiments, a highly concentrated solution of NADH at 300 mM was freshly prepared by weighing the coenzyme powder and mixing it with the corresponding crystallization buffer at 18% PEG 3350.

### Diffraction Experiments

Experiments were conducted at the SPB/SFX instrument of the European XFEL during granted proposals P3083 and P4502.^32^ During P3083, 202 pulses were delivered at 10 Hz and 9.3 keV, with an ∼3 µm spot size. The average pulse energy was 2.5 mJ with a duration of 40 fs separated by 1.77 µs, and diffraction data were collected on the AGIPD detector operating in fixed medium gain mode.^33^ During experiment 4502, 202 pulses spaced by 1.77 µs, were delivered at 10 Hz and 7 keV with a ∼3.5 µm spot size. The average pulse energy was 1.8 mJ with a duration of 40 fs, and diffraction data were again collected on the AGIPD detector operating in fixed medium gain mode.^34^ During both experiments, live hit detection was monitored via OnDA.^35^

### X-ray Diffraction Data Processing and Structure Determination

For structure determination, hits were initially identified by the Cheetah software after applying gain calibration.^36^ A crystal hit was defined to satisfy the following criteria: Using Peakfinder 8, an ADC threshold of 150, a minimum SNR of 6, and a minimum of 1 pixel per peak to identify a minimum of 10 peaks in a resolution range of 0-500 (pixels) was applied. Additionally, to correlate hits with droplets for synchronization, a Python script was used to detect instances where the hits-per-train value, from Cheetah, was non-zero. This information was then overlaid with the droplet signal for each train, generating a waterfall plot. In this visualization, green triangles were used to indicate which droplets resulted in a Cheetah-identified hit. This procedure was applied in **Figures 2a** and **3b**.

**Figure 3:**
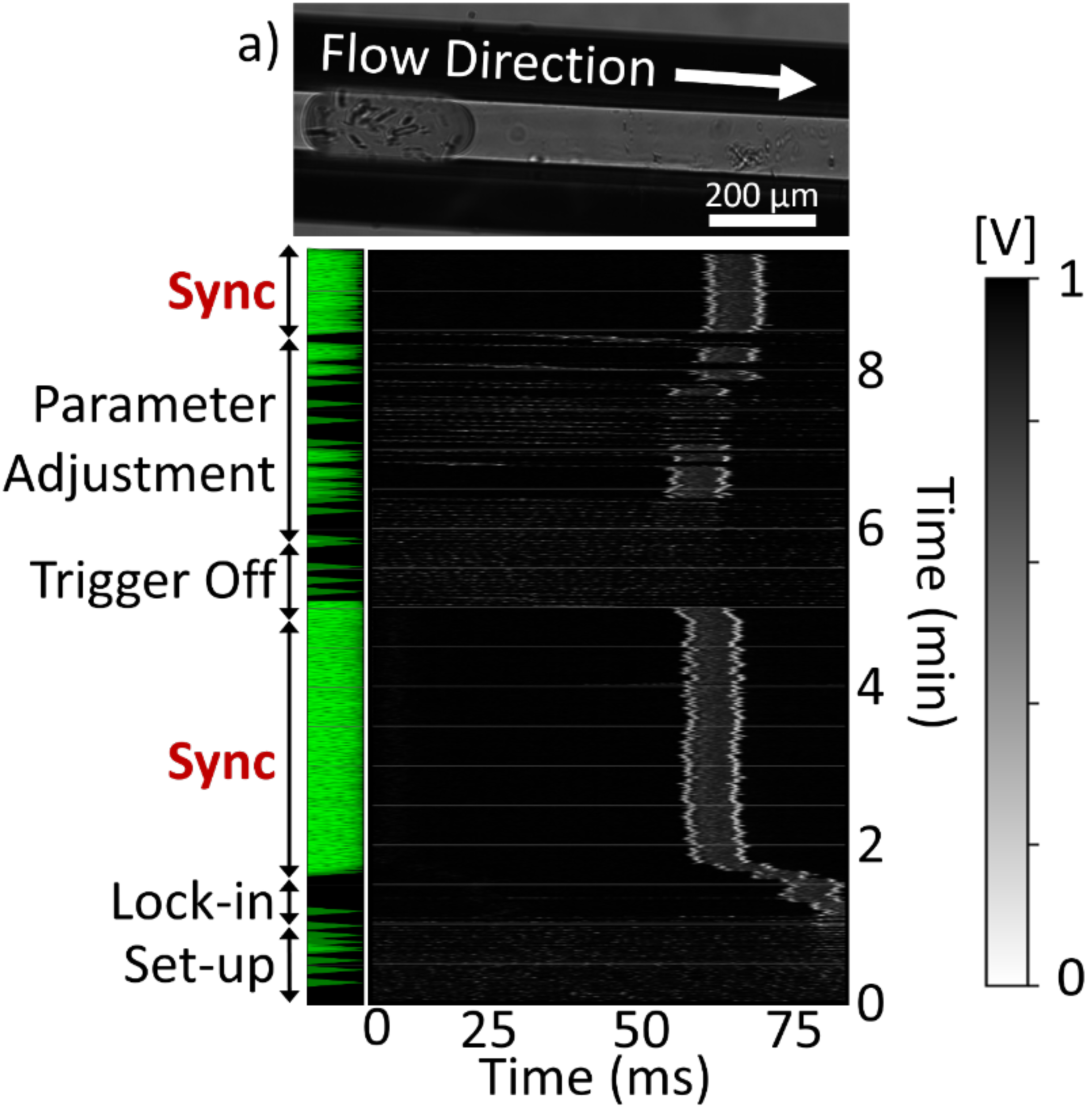
a) Image of a droplet containing crystals originating from the DG-300-Y device recorded in the SPB/SFX instrument. b) Waterfall plot of the start-up procedure conducted during P4502. The green triangles on the left indicate crystal hits.

Hits were subjected to indexing with CrystFEL v0.10.2. An additional round of peak finding with Peakfinder 8 using more robust parameters (ADC threshold of 250, a minimum SNR of 4, and a minimum of 2 pixels per peak needed to identify a minimum of 10 peaks in a resolution range of 10-800 pixels), and these peaks were subsequently supplied to indexing attempts using the algorithms Mosflm, XDS, DirAx and XGANDALF.^37–42^ Intensities were integrated applying radii of 2, 4, and 6 and merged into pointgroup mmm using the CrystFEL program partialator, applying 1 iteration of the unity model.

A representative diffraction pattern for the complex NQO1-NADH is shown in **Figure SI-1**. The final data collection and refinement statistics are listed in **Table SI-1**. During beam time P3083, two data sets, one for the free NQO1 and one for NQO1 mixed with NADH at a 305 ms reaction time point were collected. For beam time P4502, two more data sets were collected, one for the free NQO1 and one for the NQO1 mixed with NADH at the 1190 ms time point. An overview of the data collected for both beam times presented in this manuscript is shown in **Table SI-2**.

All MTZ files for phasing and refinement were generated by the CTRUNCATE program from the CCP4 software package and a fraction of 5 % reflections were included in the generated R_free_ set.^43, 44^ Initial phases of free NQO1 and NQO1 in complex with NADH were obtained by molecular replacement with MOLREP.^45^ For the free NQO1 structures, we used our recently published SFX structure (PDB 8C9J) as the search model.^22^ In the case of the dynamic structures, we used the free NQO1 structures from this study as the search model. The obtained models were refined using alternate cycles of automated refinement using non-crystallographic symmetry (NCS) with REFMAC5 and manual inspection was performed with COOT.^46, 47^ The final refined structures were validated using the Protein Data Bank (PDB) validation service prior to deposition. The atomic coordinates and structure factors have been deposited in the PDB with accession codes PDBs 9EZQ and 9EZS for the free NQO1, and 9EZR and 9EZT for NQO1 in complex with NADH at 305 ms and 1190 ms, respectively. Electron-density and POLDER maps were calculated with the MAPS tool in the PHENIX software suite.^48^ Structure figures presented in this manuscript were generated with PYMOL (version 2.4.0) (Schrödinger LLC).

## Results and Discussion

Previously, we successfully demonstrated the delivery of aqueous protein-crystal-containing droplets into the path of an XFEL both at the SLAC LCLS^22, 23^ and the EuXFEL.^24^ While synchronization was not optimized in both cases, a sample consumption reduction of up to 75% at the LCLS and up to 60% at the EuXFEL were obtained.^22–24^ Here, we further improve droplet synchronization and apply the segmented droplet injection approach for TR-SFX at the EuXFEL. For this purpose, we combined the microfluidic droplet generator with an upstream mixer to facilitate time-resolved crystallography following the mix-and-inject principle and applied it to study the reaction of NQO1 and its coenzyme NADH.

### Droplet generation with capillary coupled device at SPB/SFX

A schematic representation of the droplet injector combined with the mixing unit is shown in **Figure 1a** accompanied by the major components required for injection and synchronization as employed at the SPB/SFX instrument at the EuXFEL. Both the hardware and software elements allow the implementation of the feedback mechanism crucial for regulating droplet synchronization with X-ray pulses, a methodology previously employed at the LCLS.^23^ In contrast to the 120 Hz XFEL repetition rate at LCLS, the EuXFEL operates on a distinctive pulse structure characterized by the delivery of a train of X-ray pulses at MHz repetition rate generated every 100 ms. Within the train, 202 X-ray pulses are separated by roughly 1.8 µs in the present experiments. Given this pulse structure, our approach diverges from introducing a droplet for each pulse; instead, we aim to introduce one droplet for every pulse train. This strategy ensures that the sample droplet spans the ∼358 µs long train duration considering the apparent jet velocities and droplet volumes. Previously, characterization of the 3D-printed GDVNs with a 100 µm orifice demonstrated that jet velocities of ∼25 m/s arise with the aqueous flow rates employed in this work (18-22 µL/min) corresponding to gas mass flow rates of ∼20 mg/min.^31, 49^ Therefore, we calculated the distance traveled by a given sample volume in 358 µs to amount to 5 mm. A typical GDVN jet produced by a 3D printed nozzle is 3-10 µm in diameter.^49–51^ Further assuming a rod-like droplet shape during jetting with a radius of 5 µm (the droplet is reduced in width from the 100 µm from the GDVN liquid orifice to ∼5 µm in the jet), a minimum of 706 pL droplet volume will span the time the pulses are fired within a train. This principle is illustrated in **Figure 1a**. As we further outline below, the droplet volumes created with the droplet generator span the entire pulse train.

To employ the droplet generators during the P3083 beam time at the SPB/SFX instrument in a mix-and-inject experiment, they were coupled to upstream microfluidic mixers to enable mixing of the NQO1 crystals with the substrate NADH (see **Figure 1a&b**). The injector assembly features a Y-Mixer, bringing together the protein crystals and the substrate solution just before droplet generation. This configuration enables the execution of time-resolved serial crystallography while also conserving the sample through droplet generation.

The reaction time points explored in these two experiments are primarily influenced by the flow rates used to propel the solutions into the XFEL path. To assess the reaction time point for a specific run, the average velocity within the channel was estimated from the volumetric flow rates employed during the experiment and the length and cross-section of the fluid channels. To obtain accurate estimates, three discrete sections (A-C) were defined, where changes in flow rate and channel cross-section may occur. These sections, as illustrated in **Figure 1b**, encompass: *section A*, the length of the channel where the substrate initially encounters the protein crystal suspension with length a = 528 µm, *section B*, the length of the channel where the droplet is formed and accelerated by the oil phase with length b = 285 µm, and *section C*, the length of the capillary and nozzle before the droplet undergoes acceleration by the focusing gas within the GDVN with length c = 2 cm.

The first set of TR experiments was carried out with a mixer/droplet injector exhibiting a width, *w*, of 100 µm in *section A* and *w* of 150 µm in *section B*. This version was termed the DG250-Y-Mixer. While the droplet generator was designed to generate 10 Hz droplets, the operation of the DG250-Y-mixer at this frequency was not stable during the P3083 experiment in the SPB/SFX chamber. Nonetheless, droplet injection was continued at an average crystal flow rate (*Q_X_*) of 4.9 µL/min, substrate flow rate (*Q_S_*) of 5.0 µL/min, and oil flow rate (*Q_O_*) of 18.2 µL/min for about 40 min with the DG250-Y-Mixer. Without triggering but with droplet generation, these flow rates corresponded to a time point of about 305 ms. A more detailed discussion quantitatively assessing the mixing times and probed reaction time point is presented below. The sample flow rate during droplet injection was about a factor of 6 lower than the total flow rate (*Q_T_*) employed. Compared to continuous sample injection with a GDVN at the same *Q_T_*, approximately 83% less crystal sample was injected, resulting in the NQO1 structure at 305 ms, which will be further discussed below.

### Droplet Generator Improvements

Since our main goal is to achieve a 10 Hz droplet injection that matches the X-ray pulse pattern of the EuXFEL, factors influencing the droplet generation were investigated. First, SEM imaging was conducted to examine the critical dimensions of the droplet generator, more specifically the wall thickness and shape of the barrier separating the electrode channel from the fluidic channel where droplet generation occurs. **Figure 2a** revealed a deformation of the 5 µm thick wall that could potentially influence the effectiveness of the electrical trigger in the droplet generation region. Therefore, in subsequent versions of the droplet generation devices, the wall thickness was doubled to 10 µm. SEM imaging confirmed that the improvement in barrier thickness to 10 µm led to the elimination of the deformation (**Figure 2b**).

Additionally, further experimental characterization of the droplet generator geometry indicated that a 100 µm wide aqueous channel was suboptimal for generating 10 Hz droplets. Consequently, the sample and substrate delivery channels were widened to 150 µm matching the dimensions of the oil channel. The newly designed droplet generation devices were named DG300-Y-Mixers as illustrated in **Figure 2c-d**. Modifying the channel geometry in the droplet generation region alters the capillary number, thereby influencing the fluid interactions within the transient droplet generation regime between dripping and squeezing.^24, 52^ In addition, the taper in *section B* of the DG250-Y-Mixers was eliminated to avoid droplet break up at the junction between the 3D printed part and the capillary. Other elements of *sections A* and *C* in the DG300-Y-Mixers remained unchanged compared to the DG250-Y-Mixers.

Moreover, contact angle studies of the employed resin after printing, as summarized in **Figure SI-2,** revealed that the coating strategy of the walls of the droplet injector could be further improved. Coating the resin surfaces overnight with NOVEC 1720 (3M, St. Paul, USA) followed by thermal curing at 65°C, increased the longevity of the surface treatment when the surface is in contact with the oil phase. Additionally, these surfaces remained stable for several weeks after the treatment and curing when stored in air. **Figure SI-2a** illustrates the decay of the contact angle on a 3D printed surface over a period of up to 6 h of submersion in oil emulating the conditions of use for droplet-generating devices when transporting protein crystal-containing droplets in oil. When the surface treatment was not thermally cured, the observed decay in contact angle was more pronounced and exhibited a steeper decline compared to the surfaces that were thermally cured. This difference suggests that the longevity of the surface treatment is prolonged due to thermal curing and within a droplet generator, this extension in the lifetime of the surface treatment increases sustained functionality throughout an operational shift. The new surface treatment procedure was then implemented and applied to the DG300-Y-Mixers.

Additionally, because the preparation of an SFX beam time requires the fabrication of the mixer devices a few weeks in advance, performing a longevity study of the surface treatment was crucial (**Figure SI-2b**). The sustained contact angle on the thermally cured surfaces confirmed that the devices remained hydrophobic up to two weeks after initial surface treatment, which is advantageous for experiment preparation.

The devices were characterized thoroughly to apply the aforementioned modifications to the barrier thickness, droplet generator geometry, and surface treatment process. As illustrated in the waterfall plot in **Figure 2e**, droplets were generated using oil and NQO1 buffer at 10 Hz and exhibited long-term stability for over 2 h. These improved devices also demonstrated sufficient response to triggering, as evidenced by the rapid change in droplet phase while maintaining lock-in along the waterfall plot, see **Figure 2f**. Furthermore, the DG300-Y-Mixers created droplets that were approx. 2.3 nL in volume (see **Figure 3a** for a representative droplet image), well above the required ∼700 pL to cover the entire MHz train. Although sample savings would be optimized when utilizing smaller droplets, the DG300 geometry was further employed due to its reproducible 10 Hz droplet generation.

### TR-SFX with DG300-Y-Mixers

DG300-Y-Mixers were subsequently employed in experiment P4502 at the EuXFEL to conduct time-resolved crystallography on NQO1. The previously established feedback system was utilized to generate droplets at 10 Hz and to synchronize the droplets generated at that frequency with the EuXFEL pulse trains.^23^ The droplet generation followed a similar start-up procedure, in which substrate, crystal sample, and oil flow rates were first increased beyond the target to initiate faster delivery of solutions. After a stabilization phase of approx. 10-15 min, the droplet generation frequency reached close to the target 10 Hz, and the electrical triggering system was initiated.

Figure 3b shows a representative waterfall plot for droplet injection at 10 Hz. In this representation, the droplet detector trace is divided into 100 ms sections according to the XFEL-reference at 10 Hz and each trace is aligned in a vertical stack with the color representing the amplitude of the signal. As the flow rates stabilized at *Q_X_* = 1.00 µL/min, *Q_S_* = 0.75 µL/min, and *Q_O_* = 18.5 µL/min, droplets were created near the 10 Hz target frequency. The external trigger was then activated with a duration of 4 ms, and an amplitude of 180 V. Droplets were stabilized just before 2 min maintaining stability for approx. 3 min without any system disturbance. Green triangles on the left, indicating crystal hits, confirm synchronization with the EuXFEL pulse structure for these 3 min, with 1631 diffraction patterns recorded at a hit-rate of 0.4%.

Following this stack of synchronized droplets, the trigger was turned off after 5 min, causing the droplets to immediately fluctuate in frequency and phase. A minute later, the trigger was re-activated with the same duration and delay, but with a reduced amplitude of 40 V. Although the droplet signal eventually locked-in about 30 seconds later, it did not remain stable, indicating that the reduced amplitude was not sufficient to stabilize the droplet-generation frequency lock-in. Similarly, at ∼7.5 min, the amplitude was increased to 110 V, resulting in more frequent but still unstable lock-ins. Finally, at ∼8.5 min, the amplitude was increased back to 180 V causing the droplets to immediately lock-in and synchronize as observed initially. This second instance of synchronization demonstrates the reproducibility of the synchronization process once droplets are generated at 10 Hz and the appropriate triggering conditions are applied. As illustrated in **Figure SI-3**, 10 Hz droplets could be generated through electrical triggering for durations of up to one hour as indicated through waterfall plots of 12 consecutive runs. These results validate the synchronization of the triggered droplet injection approach for the first time at 10 Hz at the SPB/SFX instrument. The long-term stability of the droplet synchronization was also explored.

### Time Points for TR-SFX with Y-Mixers

With the flow rate parameters described above to achieve synchronized droplet injection at 10 Hz, we will now discuss the time points probed in the reaction of NQO1 with NADH, as well as the mixing times achieved to initiate this reaction. The two DG-Y-Mixer devices can be broken down into three sections as previously outlined: *section A*, where the substrate and crystal stream first meet and mixing is initiated by diffusion in the convective flow, *section B*, where the droplet is formed and additional mixing may occur in the developing droplet, and *section C*, the path of the droplet through the remaining device prior to injection by the GDVN. The sum of the three sections and corresponding flow rates determines the average time point of the reaction probed, *t_R_*, illustrated in equation 1:

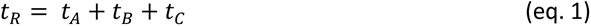

In the DG250-Y-Mixer, the flow rates of *Q_X_* = 4.9 µL/min and *Q_S_*= 5.0 µL/min, were used during the mixing experiment. Based on the device geometry and flow rates, the sample spends an average time, *t_A_*, of 32 ms in this first section. Once the droplet is formed, with *Q_O_* of 18.2 µL/min, the total flow rate *Q_T_* amounted to 28.1 µL/min leading to *t_B_* and *t_C_* of 14 and 259 ms respectively, and *t_R_* of 305 ms.

This contrasts with the DG300-Y-Mixer, which was operated at lower aqueous flow rates to generate 10 Hz droplets. In the DG300-Y-Mixer, the longer residence time in *section A* resulted in *t_R_* of 1190 ms before irradiation with the XFEL pulses. *Q_X_* = 0.5 µL/min, *Q_S_* = 0.4 µL/min, and *Q_O_* = 18.3 µL/min resulted in *t_A_*, *t_B_* and *t_C_* as summarized in **Table 1**.

**Table 1:** Average Residence Times for Solutions in Droplet Generators.

| Droplet Generator | $t_A$ (ms) | $t_B$ (ms) | $t_C$ (ms) | $t_R$ (ms) |
| --- | --- | --- | --- | --- |
| DG250-Y-Mixer | 32 | 14 | 259 | 305 |
| DG300-Y-Mixer | 792 | 20 | 378 | 1190 |

We considered that the primary contribution to the spread of *t_R_* was caused by the diffusion-based mixing in *section A* and the mixing occurring in the droplet once formed in *section B*. Once incorporated in droplets, crystals are transported at the same velocity, thus the time variation in *section C* is negligible. To quantitatively determine the spread of *t_R_* due to the different mixing regimes, the beginning of the reaction was assumed to occur when the concentration of the protein in the crystal and the substrate concentration were equivalent. Furthermore, to estimate the mixing rate of the substrate into the crystal stream, a convection-diffusion model using finite element modeling was employed. This was accomplished with a 2D-model that includes a channel section corresponding to *section A*, flow rates as employed for the experiments with the two time points, and with a reported diffusion coefficient for NADH of 6.7 ∗ 10^-6^ *cm*^2^*s*^-1^ ^53^. The model is outlined in the methods section and **Table SI-3**.

The concentration distribution arising from the convection at the employed flow rates and diffusion of NADH were evaluated at two different positions within *section A;* at partition x (where the two streams meet) and partition y (at the end of the channel, just before the droplet will be formed), schematically represented in Figure 4a. At these different partitions, the projection of the concentration profiles are depicted in **Figure SI-4a and b** for the DG300-Y-Mixer and DG250-Y-Mixer, respectively, were point *n* corresponds to the point at which the substrate concentration and the protein concentration in the crystal are equimolar, and where the reaction will initiate.

**Figure 4:**
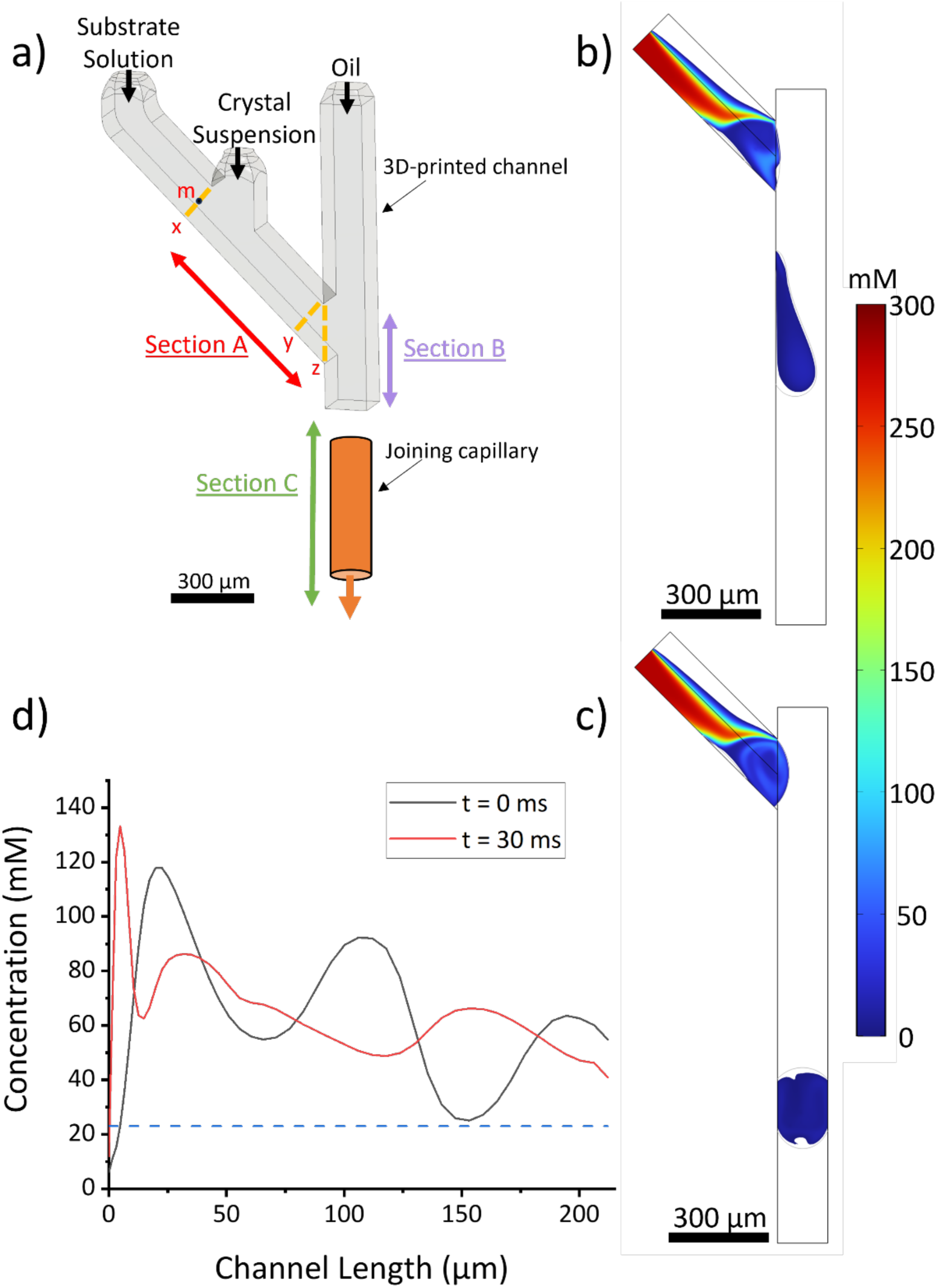
(a) Schematic representation of the mixing droplet generation device, divided in the partitions used to evaluate the diffusion of the substrate molecule into the crystal sample stream. (b) The result of the 2D computational model overlaying the droplet volume interface with the substrate concentration for the DG-300-Y device at the point when the previous droplet just broke off from the sample channel (t=0) and c) as in b), but at 30 ms after droplet break off. d) concentration of substrate along line z (as indicated in a) where sample and oil stream meet at t=0 ms and t=30 ms.

Where the two solutions (crystal and substrate) first meet in the center of the channel in *section A* (at point *m* in Figure 4a) signifies the earliest instance at which the reaction can initiate, representing the longest reaction time. Due to a parabolic flow profile, the mixing of the substrate with crystals was not homogeneous so the crystals that are closer to the channel wall flowed at a lower velocity and, thus, mixed with the substrate later. Therefore, we determined both the slowest and fastest velocities of the aqueous flow and the resulting difference in travel time contributing to the range in the calculated reaction time. From this model in the DG300-Y-Mixer, we noticed that the substrate concentration in the solution carrying the crystal sample at the end of *section A* had only reached a 1:1 mixing ratio 36 µm from the center of the channel (or, correspondingly, 39 µm from the channel wall) (**Figure SI-4a**). This indicated that only about 50% of the sample had begun to react, therefore, comparing the linear flow velocity at this position with that in the center of the channel, resulted in a 55.4 ms difference for the fastest and slowest flow velocity in the channel in which the substrate concentrations were large enough to initiate binding.

In the DG250-Y-Mixer, since the solutions flowed significantly faster within *section A*, the substrate reached an equivalent concentration to the crystal solution about 8 µm from the center of the channel at the end of *section A* at point *n* (see also **Figure SI-4b**). Due to the lowered residence time in *section A*, about 85% of the solution within the channel remains unmixed before the droplets are generated. Therefore, the mixing time variation in this region resulted in 0.15 ms as a result of the difference between the fastest and slowest flow velocity in the channel in which the substrate concentrations were large enough to initiate binding.

In *section B*, the two liquid streams entered the continuous flow channel forming droplets embedded in the oil phase that, in turn, induced a spontaneous circulation mixing effect that further enhanced mixing. Therefore, to simulate this mixing process, the formation of the droplets in the continuous oil channel was studied with a 2D model of the droplet device geometry. A laminar two-phase flow model employing the level set interface method was combined with the transport of diluted species module to simulate the droplet formation and the diffusion of the substrate molecules. The two phases considered for simulation included the fluorinated oil mixture and NQO1 mother liquor (see methods section and **Table SI-3** for details). The model was built as previously described,^31^ however, the parameters were adapted to the experimentally used liquids and device geometries.

Two representative snapshots from this time-dependent model are depicted in Figure 4b and c for the DG300-Y-Mixer (1190 ms time point). The first snapshot (Fig. 4b) corresponds to the frame after a droplet has just broken off the intersection. Closer analysis of the concentration distribution along the line separating the sample and oil channel, as plotted in Figure 4d, indicates that mixing occurred, as the substrate concentration surmounts 23 mM. This specific concentration corresponds to a 1:1 mixture of protein to substrate based on the estimated concentration of NQO1 in the crystal. Interestingly, the flow profile dictated by the differences in flow rates in the two phases lead to an additional mixing effect (observed from the concentration distribution at the Y-intersection in Figure 4b), which almost instantaneously increases the substrate concentration to > 23mM at the position where the droplet is formed. The second snapshot in Figure 4c (30 ms after the start of droplet formation), confirms complete mixing, which is also reflected by the concentration profile in Figure 4d. We thus conclude, that mixing in the droplets is instantaneous, and no further spread in the mixing time is introduced through droplet formation.

The same mixing characteristics were also found for the 305 ms time point in the DG250-Y mixer. Analysis along the center line, as evidenced in **Figure SI-7**, confirms that the time spread variation due to droplet formation is negligible as the substrated concentration surmounts 23mM. Thus, the total time spread of *t_R_* calculated for the DG300-Y-Mixer remained at 55 ms (contribution from section A) and at 0.15 ms for the DG250-Y-Mixer. Lastly, we compare the variations in mixing times with the diffusion of the substrate into the protein crystals. For shoe-box shaped crystals with dimension of 10 x 20 x 30 µm^3^, diffusion times of ∼15 ms were reported. ^54^ Thus, accounting for similar time scales in our case, the time spread for both time-resolved experiments is below 6% of the reaction time point probed for each case.

Based on the above presented analysis it is evident that the length of *section A* poses a limitation when attempting to investigate reactions at faster time points than reported here, specifically when operating the droplet generator at 10 Hz, where substrate and sample flow rates are small. Consequently, future efforts will involve the design and evaluation of mixing devices where the mixing occurs shortly before the droplet generation region. This approach aims to not only minimize overall reaction time but also restrict the time spread introduced by diffusive mixing of the samples. For example, in designs with channel dimensions similar to those for the DG300, but where the length of *section A* would be reduced from 528 µm to 100 µm, the time spread can be reduced to 11 ms.

### Sample Consumption

For the 305 ms and 1190 ms time points explored in our experiments, we further provide an assessment of sample consumption in the droplet injection mode, as shown in **Table 2**. While droplet generation was sustained throughout the experiment at the EuXFEL for several hours, the data presented in this table only includes runs where droplet generation was optimized for diffraction collection for the two specific time points targeted.

**Table 2:** Time points analyzed and sample consumption during P3083 and P4502.

| $t_R$<br>(ms) | Data<br>Collection<br>Time (min) | Indexed<br>patterns | Patterns/$\mu\text{L}$ | Protein<br>Concentration<br>(mg/mL) | Protein<br>Consumption<br>(mg) | Resolution<br>($\text{\AA}$) |
| --- | --- | --- | --- | --- | --- | --- |
| **305** | 38 | 18,794 | 101 | 18 | 3.2 | 2.7 |
| **1190** | 85.5 | 10,992 | 257 | 25.5 | 1.7 | 2.5 |

For the 305 ms time point, 3.2 mg of protein sample was consumed which is notably much less than that required from continuous injection with a GDVN, estimated to be 18.5 mg at the same total flow rate and time. This equates to nearly 6-fold sample savings despite the lack of droplet synchronization. At the ideal 10 Hz frequency with the DG300-Y-Mixer, droplet injection was recorded for about 1.5 h, generating nearly 11,000 patterns, resulting in 257 patterns per µL of sample injected. This was about 2.5 times more efficient than for the 305 ms time point. Furthermore, for the 10 Hz scenario where the longer time point (1190 ms) in the reaction of NQO1 with NADH was probed, more than 60 mg of protein would have been consumed if a continuous GDVN were employed with a similar flow rate. Thus, droplet injection conserved 97 % of the sample compared to a continuous injection demonstrating the advantage of using the droplet injector as a sample delivery device for mix-and-inject TR crystallography.

### Novel structural and dynamics insights into the reductive half-reaction of NQO1 with NADH

Efficient reaction initiation in crystals requires fast incorporation of ligands to the active sites. To this end, in a *proof-of-principle* experiment by our collaborative group, we demonstrated that NQO1 microcrystals are suitable for time-resolved experiments through confirmed binding of the coenzyme NADH with the crystal slurry after incubating for 1h before analysis.^27^ To further investigate the molecular determinants of the catalytic mechanism of NQO1, here we present novel time-resolved SFX data of the reductive half-reaction of NQO1 with NADH at two mixing time points (305 and 1190 ms). For comparison, we have also determined the crystal structures of NQO1 in its free form. It is important to note that, since we are focusing on the segmented droplet injection method facilitating TR-SFX, a more comprehensive analysis of the structural data presented here, along with computational and biophysical studies, will be forthcoming.

NQO1 microcrystals belonged to the space group P2_1_2_1_2_1_ with two homodimers in the asymmetric unit (Figure 5a) related by a non-crystallographic two-fold axis of symmetry with a typical folding characteristic .^55^ All data collection, processing, and refinement parameters and statistics are given in **Table SI-1**. As reported by our group in two recent studies^22, 27^ there exists a high conformation heterogeneity in the two catalytic sites of NQO1 that agrees with the negative cooperative previously reported.^56–59^ To see if this is also observed in the free NQO1 structures reported here, we carried out a structural comparison of the free NQO1 structures from P3083 and P4502 experiments with each other and with other free NQO1 structures determined at room temperature by serial crystallography.^22, 27^ As illustrated in **Figure SI-5** and in good agreement with what we have previously reported,^22, 27^ residues at the catalytic site show a high flexibility. Among the highest flexibility is observed for residues Asn64, Gln66, Tyr126, Arg200 and Asn233, the most relevant for Tyr128 and Phe232, which, for some of the homodimers, were modeled in various conformations. The Tyr126 and Tyr128 residues, that gate the catalytic pocket of NQO1, are strictly conserved and have been shown to be the key players in the function of NQO1.^60, 61^

**Figure 5:**
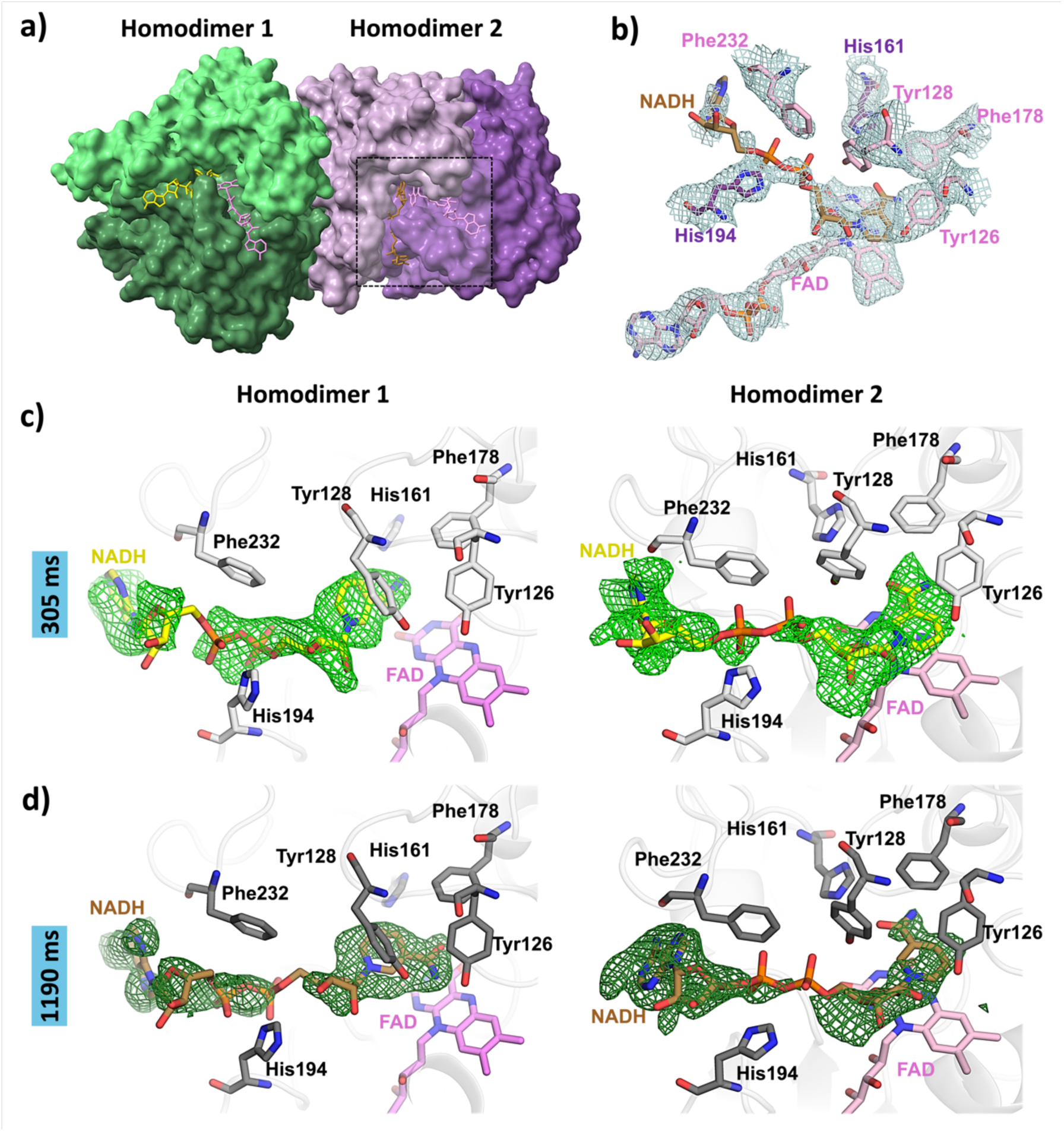
Time-resolved structures of NQO1 with NADH at 305 and 1190ms. **a)** Surface representation of the two homodimers of NQO1 found in the asymmetric unit. The individual monomers are highlighted in light green (chain A), dark green (chain B), light pink (chain C), and purple (chain D). The NADH molecules bound to chain B and chain D, are represented as sticks in yellow and golden, respectively. The cofactor FADs are shown as pink sticks. **b)** Electron density maps 2mFo-DFc contoured at 1 σ and stick representation of the FAD, NAD, and the residues in the catalytic site highlighted for the dash boxed panel in a). **c)** POLDER maps contoured at 3 σ of the NADH molecule bound to homodimer 1 (left) and homodimer 2 (right) at 305 ms time point. **d)** POLDER maps contoured at 3 σ of the NAD molecule bound to homodimer 1 (left) and homodimer 2 (right) at 1190 ms time point.

In contrast to what has been reported by us for the structure of NQO1 with NADH at *t_R_* = 1 hour, no significant difference in the unit cell dimensions between the free and mixed proteins was observed.^27^ This could be due to the fact that the reduction in the plasticity of NQO1 reported upon binding of NADH takes longer than the time points presented in this study. The resulting experimental maps were of excellent quality revealing, besides the presence of FADs, many water molecules **Table SI-1**. The high quality of the NQO1 structure can be assessed from the 2mFo-DFc electron density maps shown for the catalytic site residues and the cofactor FAD (Figure 5b). Further evaluation of the quality of the free NQO1 structures versus those with NADH bound was carried out by comparing crystal structures recently reported at room temperature (PDB entries 8C9J and 8RFN for the free NQO1 and PDB entry 8RFN or the NQO1 in complex with NADH).^22, 27^ Overall, all NQO1 structures aligned very well with each other, with average root-mean-square deviation (RMSD) values of 0.294 Å for the Cα atoms. The global RMSD of 0.590 Å suggests slightly higher structural differences when the whole protein molecule is considered, mainly due to mismatch from flexible loops as well as solvent-exposed regions as expected.

POLDER maps of the NADH molecules in the final model of the complex at 305 and 1190 ms are shown in Figures 5c **and d**, confirming the lack of model bias and, thus, their binding to the enzyme. Strong electron density is already observed at 305 ms, indicating fast ligand binding even under a more viscous mother liquor condition and the large size of the NADH molecule. These results demonstrate that ligand binding occurs within the millisecond time domain. At the 1190 ms time point, the intensity of the electron density does not change much compared to that of 305 ms. This could be explained by the extremely high flexibility of the NADH molecule at the catalytic site of NQO1.^27^ It is important to note that, at both times in the reaction, the NADH molecules are not bound to the same homodimer, but one of them is bound to one active site of one of the homodimers and the other one is bound to one of the active sites of the other homodimer in the asymmetric unit (Figure 5a). Such observation is in agreement with the negative cooperativity previously reported from multiple biophysical and biochemical studies in solution.^56, 57, 59, 62^Furthermore, the exact disposition of both NADH molecules within each time point differs from one another and is also different from the position and orientation of the NADH molecule in our recently published static synchrotron structure of NQO1 with NADH (**Figure SI-6**).^27^

Because NQO1 is involved in the detoxification processes within cells, particularly in the reduction of quinones, the negative cooperativity or binding inhomogeneity observed in NQO1, would allow NQO1 to efficiently handle varying concentrations of quinones within cells. When one quinone substrate molecule binds to the active site of NQO1, it induces a conformational change in the enzyme that reduces the affinity for additional quinone substrate molecules. Overall, negative cooperativity in NQO1 might be relevant because it allows the enzyme to regulate its activity in response to changes in substrate concentration, contributing to cellular detoxification processes and maintaining cellular homeostasis.

## Conclusion

We have demonstrated segmented droplet generation as an effective method to facilitate MISC TR-SFX at the SPB/SFX instrument at the EuXFEL for the first time. Our method differs from drop-on-demand approaches, as a segmented stream of oil phase separated by aqueous crystal-laden droplets are continuously injected and thus maintains all characteristics of liquid jet injection with nozzles while it is compatible with MHz repetition rates. We demonstrate the synchronization with the pulse trains of the EuXFEL repeating at 10 Hz for the first time and additionally couple a mixer prior to droplet generation to investigate the reaction of NQO1 with its substrate NADH at 305 ms and 1190 ms. With droplets generated at 10 Hz frequency, we successfully solve the structure of NQO1 with the substrate NADH bound at a resolution of 2.5 Å. Compared to a continuous injection with a GDVN at the same sample flow rate, 97% of the sample could be conserved.

We have further demonstrated TR-SFX for the reaction of NQO1 with NADH at the 305 ms reaction time point. Although droplet generation was not stable at a defined frequency, this approach still saved up to 85% sample and thus represents a pathway towards faster time points with minimal geometric changes in the droplet generator. Future improvements on further optimizing this droplet synchronization approach for TR-SFX include the minimization of the spread in the reaction time points, which can be achieved by design changes such as placing the convection-diffusion-based mixer closer to the droplet generation section and additionally taking advantage of fast turbulent mixing in the droplets eliminating the need for the upstream mix-and-inject approach.

We further report on the first structural studies of the NQO1 reaction with NADH in a TR-SFX experiment, indicating NADH binding to the NQO1 homodimer. A detailed structural analysis of the interaction of NQO1 with NADH is beyond the scope of this manuscript. However, the structural information reported here provides new insights into the catalytic function of NQO1 and will pave the way towards deciphering the reaction mechanism of this important therapeutic enzyme. A comprehensive structural analysis study coupled with biophysical and computational methods elucidating details of the reaction mechanism will be forthcoming.

## Supporting information

Complete Supplementary Material

## Acknowledgments

We acknowledge the European XFEL in Schenefeld, Germany, for granting experimental time for the proposals P3083 and P4502 as well as off-shift collaborative efforts on the SPB/SFX (Single Particles, Clusters, and Biomolecules, and Serial Femtosecond Crystallography) instrument and we would like to thank the instrument operations group as well as the faculty and staff for their assistance. We also acknowledge the use of the Auriga SEM within the Eyring Materials Center at Arizona State University supported in part by NNCI-ECCS-1542160.

The team further acknowledges funding from the National Science Foundation through the BioXFEL Science and Technology Center (agreement #1231306), as well as under midscale RI-2 award No. 2153503 and DBI award No. 1943448.

The European Union NextGenerationEU/PRTR (Grant number CNS2022-135713) and the Ayuda de Atracción y Retención de Talento from the Community of Madrid (Grant number 2019-T1/BMD-15552), the Spanish State Research Agency and FEDER (MCIN/AEI-FEDER, Grant PID2022-136369NB-I00) and the Government of Aragón-FEDER (Grant E35_23R) are also acknowledged for financial support.

