## Supplementary material for "Minimized Sample Consumption for Time-Resolved Serial Crystallography Applied to the Redox Cycle of Human NQO1": Complete Supplementary Material

**Table SI-1: Data Collection and Refinement Statistics (values for the outer shell in parentheses)**

| | Free NQO1 (P3083) | Free NQO1 (P4502) | NQO1-NADH (305 ms) | NQO1-NADH (1190 ms) |
|---|---|---|---|---|
| *Data collection statistics* | | | | |
| Data collection time (min) | 66.5 | 191.5 | 38 | 85.5 |
| Wavelength (Å) | 1.3332 | 1.7712 | 1.3332 | 1.7712 |
| Detector | AGIPD 1 Mpx | AGIPD 1 Mpx | AGIPD 1 Mpx | AGIPD 1 Mpx |
| Space group | $P2_12_12_1$ | $P2_12_12_1$ | $P2_12_12_1$ | $P2_12_12_1$ |
| a, b, c (Å) | 61.6, 107.6, 198.6 | 61.5, 107.8, 198.1 | 61.6, 107.7, 198.6 | 61.5, 107.8, 198.1 |
| α, β, γ (°) | 90, 90, 90 | 90, 90, 90 | 90, 90, 90 | 90, 90, 90 |
| Resolution range (Å) | 27.0-2.5 (2.56-2.50) | 26.7-2.30 (2.38-2.30) | 25.6-2.5 (2.57-2.51) | 26.4-2.5 (2.59-2.50) |
| Completeness (%) | 100 (100) | 100 (100) | 100 (100) | 100 (100) |
| CC* (%) | 98.55 (89.13) | 98.59 (71.56) | 98.72 (82.0) | 97.86 (54.64) |
| $CC_{1/2}$ (%) | 94.39 (65.88) | 94.55 (34.41) | 95.06 (50.64) | 91.86 (17.5) |
| Multiplicity | 489 (347) | 237 (142) | 287 (204) | 115 (84) |
| $R_{split}$ | 20.4 (69.5) | 22.5 (126.8) | 22.0 (127.1) | 28.5 (189.8) |
| Avg. I/σ (I) | 4.3 (1.2) | 3.5 (0.6) | 3.7 (0.7) | 2.8 (0.5) |
| *Refinement Statistics* | | | | |
| Resolution range (Å) | 27.0-2.5 (2.56-2.50) | 26.7-2-30 (2.36-2.30) | 25.6-2.5 (2.57-2.51) | 26.4-2.5 (2.56-2.50) |
| No. of reflections, working set | 44,040 | 56,465 | 42,064 | 43,444 |
| No. of reflections, test set | 2,350 | 2,939 | 2,176 | 2,326 |
| $R_{work}/R_{free}$ (%) | 23.7 / 19.4 | 20.0 / 24.9 | 20.6 / 25.5 | 19.9 / 24.4 |
| No. of non-H atoms | | | | |

| | | | | |
|---|---|---|---|---|
| Protein | 8606 | 8764 | 8725 | 8691 |
| Water | 279 | 579 | 224 | 247 |
| FAD/NADH | 208/0 | 208/0 | 208/86 | 208/86 |
| Others | 0 | 4 | 0 | 0 |
| R.m.s. deviations | | | | |
| Bond length (Å) | 0.006 | 0.006 | 0.005 | 0.006 |
| Bond angles (°) | 0.001 | 0.001 | 0.001 | 0.001 |
| Average *B* factors (Å$^2$) | 46.99 | 41.10 | 56.23 | 28.11 |
| Ramachandran plot | | | | |
| Favored (%) | 97 | 98 | 98 | 97 |
| Allowed (%) | 3 | 2 | 2 | 3 |
| Outliers (%) | 0 | 0 | 0 | 0 |
| PDB code | 9EZQ | 9EZS | 9EZR | 9EZT |

**Table SI-2: Data Acquisition During P3083 and P4502**

| Beam Time | Dataset | Frames | Hits | Indexed Patterns | Lattices |
|---|---|---|---|---|---|
| **P3083** | Free NQO1 | 8,411,372 | 44,508 | 35,329 | 38,226 |
| | 305 ms | 4,422,184 | 24,631 | 18,794 | 19,815 |
| **P4502** | Free NQO1 | 14,656,518 | 40,168 | 28,877 | 34,367 |
| | 1190 ms | 9,490,617 | 15,268 | 10,992 | 12,903 |

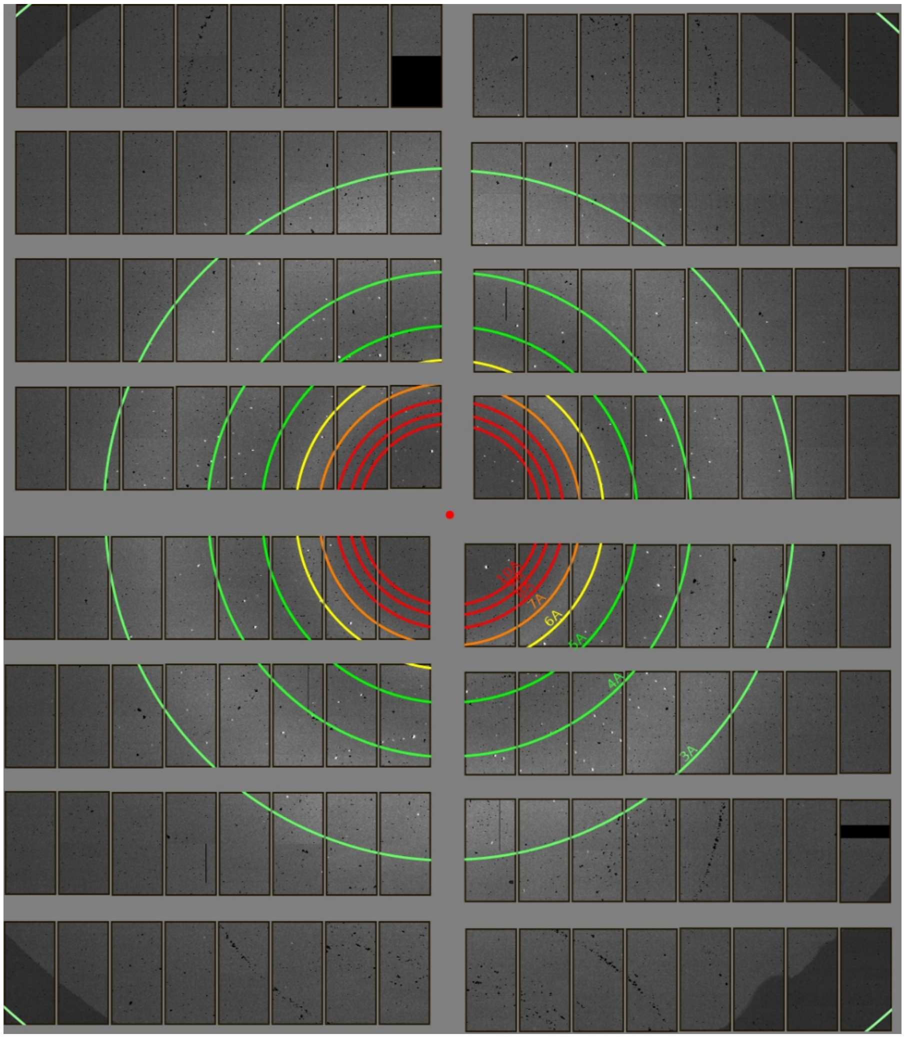

**Figure SI-1:** A representative diffraction pattern obtained for the 1190ms time point.

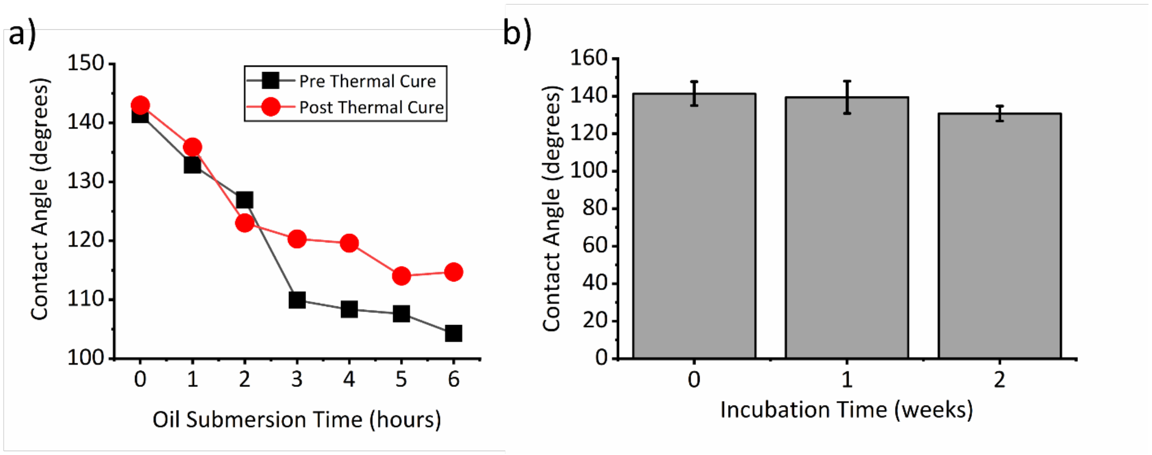

***Figure SI-2:*** *Contact angle characterization following the previously established procedures for the boundary created between oil, water, and 3D-printed substrate.*[1] ***a)*** *Contact angle decay dependent on the use of thermally cured and not thermally cured surface treatment.* ***b)*** *Contact angle decay after thermal treatment when incubated in air at room temperature.*

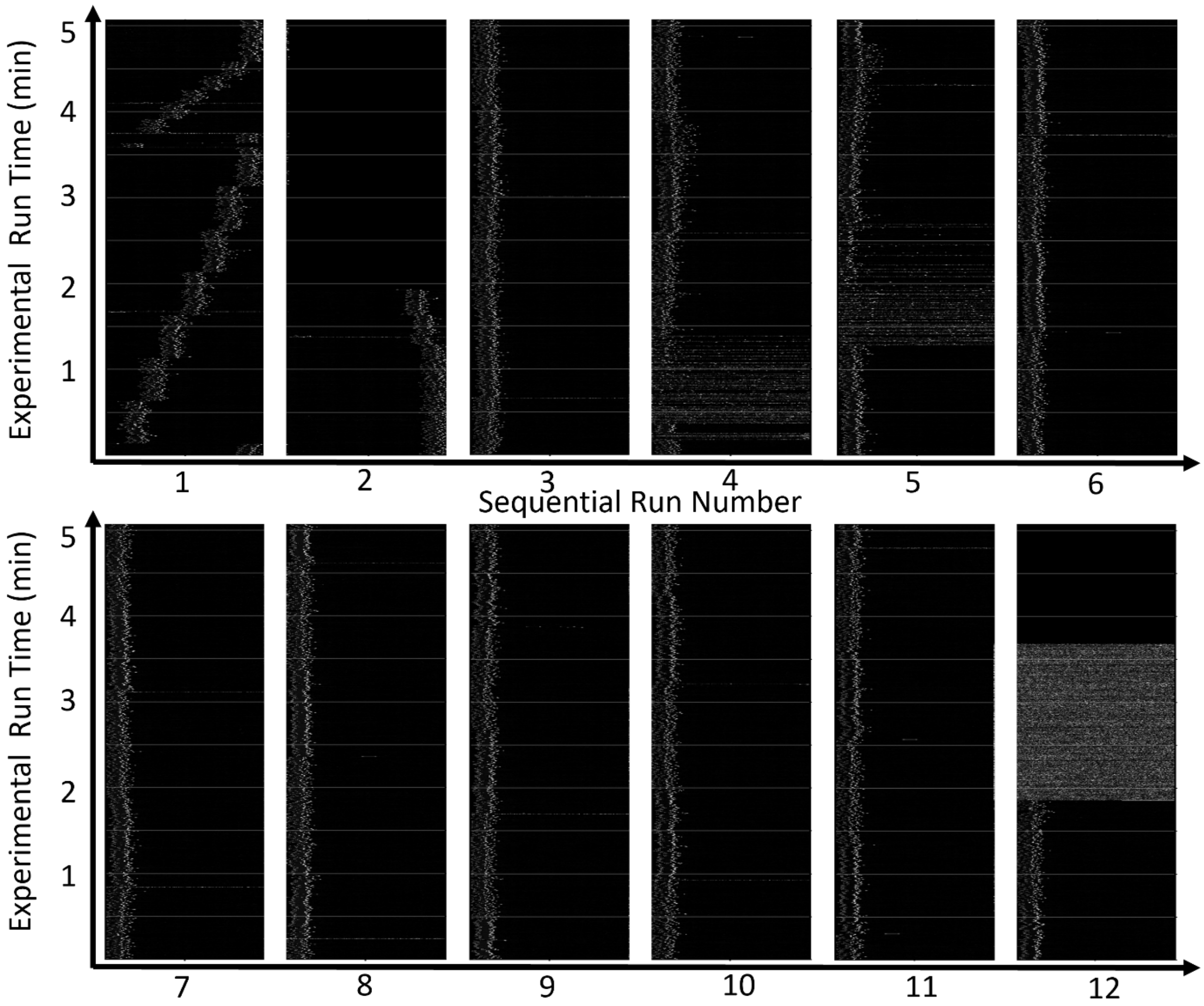

***Figure SI-3:*** *Waterfall plot collected during P4502 during droplet delivery of mixed NQO1 crystals over 12 runs equating to one hour of continuous data collection. The droplets are initiated and locked-in within the first minute. Subsequently, the droplets are manually scanned along the period of the XFEL in the following 4 min until an optimal delay is obtained. Finally, the droplets remained locked-in at the same delay for 9.5 runs (50 mins) before the droplets were intentionally stopped by turning off the liquid flow and disturbing the droplets around the 2-minute mark of the final run.*

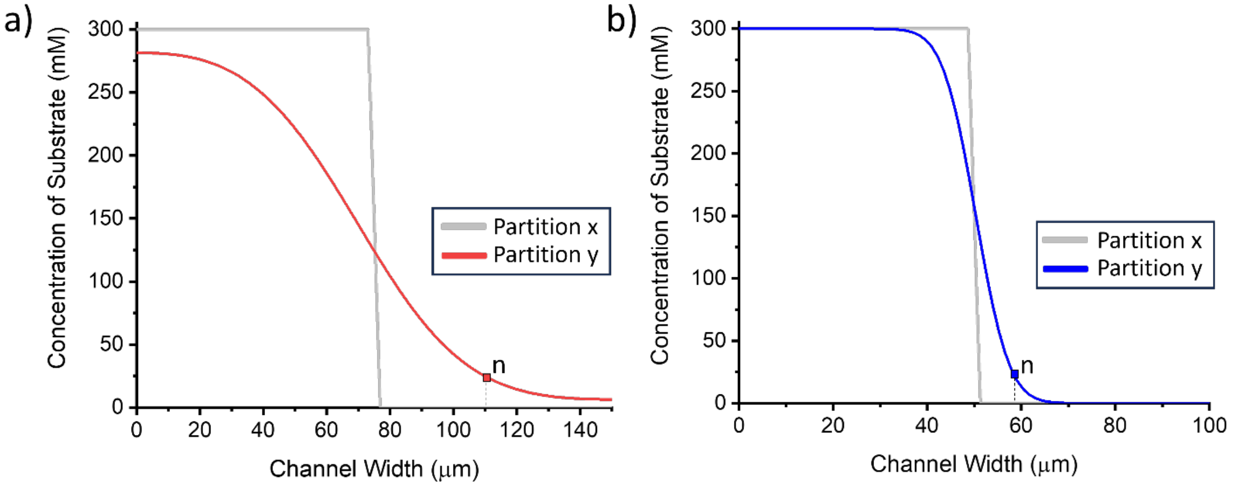

***Figure SI-4:*** *Numerical Simulation results for diffusion of substrate molecules in solution across channel width for **a)** the DG300-Y-Mixer with concentration sampled at the beginning (x) and end (y) of the section A where the initiation of the reaction is initiated at point n (where 1:1 substrate and protein concentration occurs). **b)** Concentration profile for the DG250-Y-Mixer at the beginning (x) and end (y) of* section A.

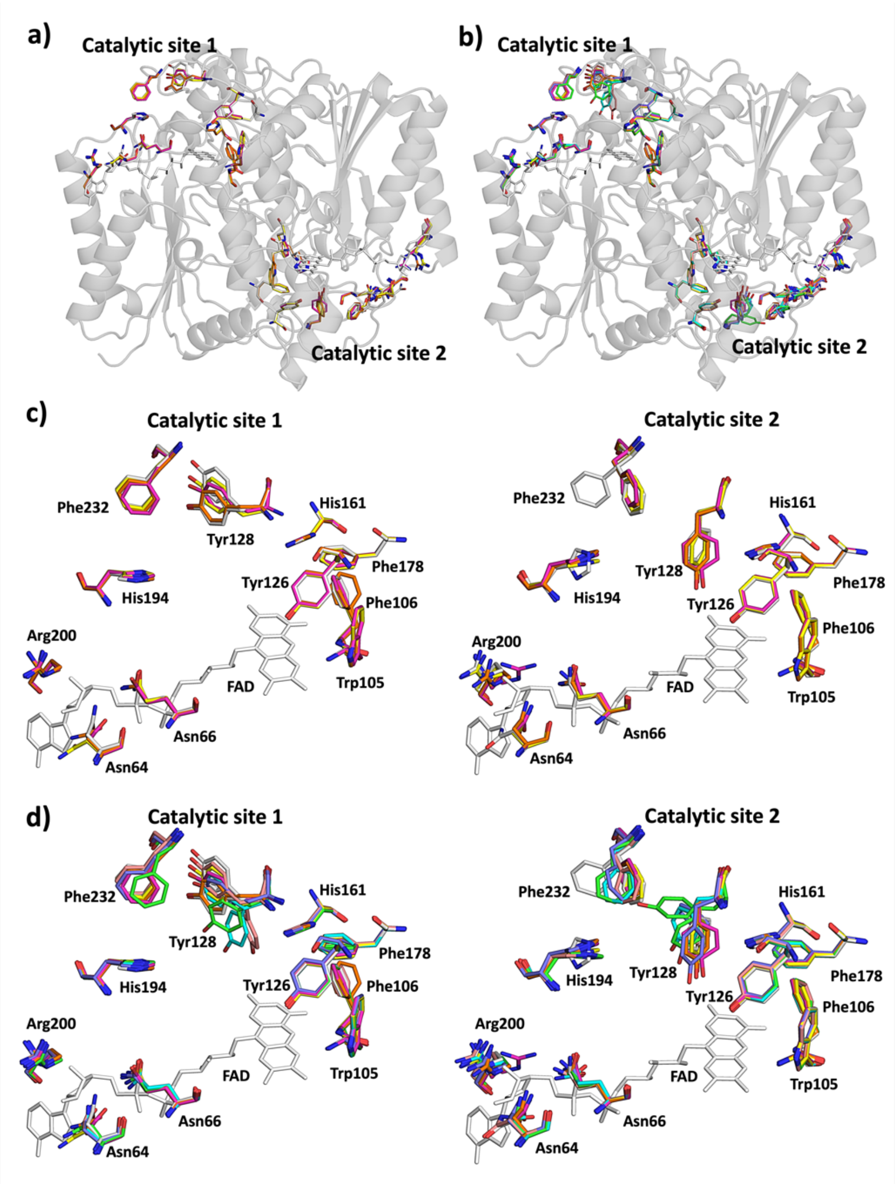

***Figure SI-5.*** *Structural comparison of the free NQO1 structures. a) Superposition of the two homodimers of the free NQO1 structures reported in this study (P3083 (homodimer 1 (magenta) and homodimer 2 (yellow) and P4502 (homodimer 1 (orange) and homodimer 2 (white)). For clarity, only one NQO1 homodimer is shown as a cartoon representation in dark grey and the residues and the FAD in the catalytic sites of all homodimers are shown as sticks. b) Superposition of the two homodimers of the free NQO1 structures reported in this study shown as is a), and those from other free NQO1 structures reported using serial crystallography (homodimer 1 (green) and homodimer 2 (cyan) for PDB 8C9J (Doppler et al. 2023); homodimer 1 (violet) and homodimer 2 (salmon) for PDB 8RFM (Grieco et al. 2024). For clarity, only one NQO1 homodimer is shown as a cartoon representation in dark grey and the residues and the FAD in the catalytic sites of all homodimers are shown as sticks. c) A closer view of the structural differences observed in the catalytic sites 1 and 2 shown in a). d) A closer view of the structural differences observed in the catalytic sites 1 and 2 is shown in b).*

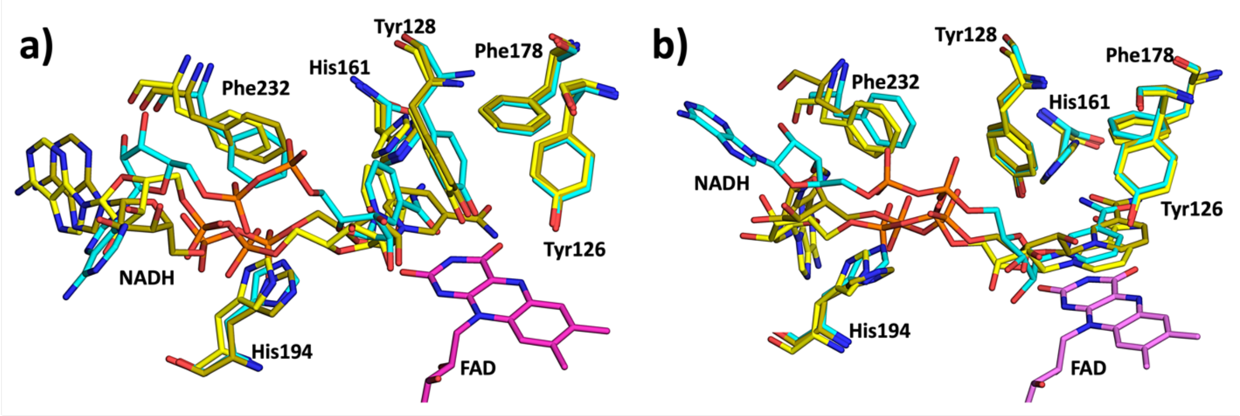

***Figure SI-6.*** *Structural comparison of the catalytic sites.* ***a)*** *Stick representation of the catalytic sites of the homodimer 1 at 305 ms (yellow), 1190 ms (dark yellow), and t = infinite (cyan).* ***b)*** *Stick representation of the catalytic sites of the homodimer 2 at 305 ms (yellow), 1190 ms (dark yellow), and t = infinite (cyan)*

### Supplementary Figures for Numerical Modeling

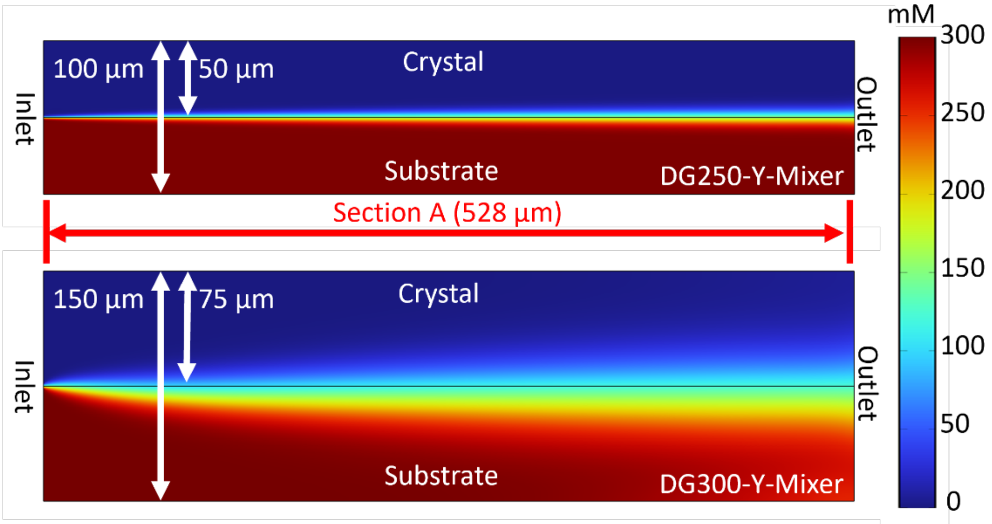

***Figure SI-6:*** *2D channel dimensions used for convection-diffusion simulation for the DG250-Y-Mixer (top) and DG300-Y-Mixer (bottom). The substrate begins in the lower half of the channel (red) and diffuses into the crystal stream (blue) at the upper half of the channel. Color gradient represents the concentration of substrate in mM.*

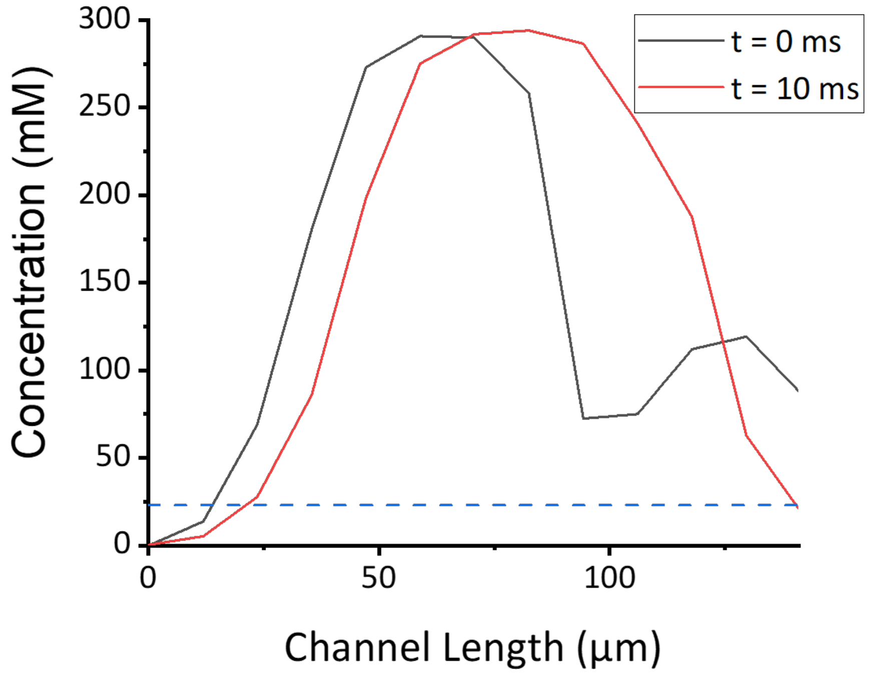

**Figure SI-7:** Concentration of substrate along line z (as indicated in **Figure 4a**) where sample and oil streams meet at t=0 ms and t=10 ms for the DG250-Y-Mixer.

**Table SI-3.** Boundary conditions, relevant equations, and modified parameters used for simulations (unless detailed, the parameters remain unchanged from our previous study).[1]

| **Physics** | **Boundary Conditions and Modified Parameters** |
| --- | --- |
| Laminar Flow | Surface Domain:<br>$0 = -\nabla p + \mu \nabla^2 \boldsymbol{u}$<br>$\rho \nabla \cdot (\boldsymbol{u}) = \boldsymbol{0}$<br>**Wall:**<br>$u = 0$ (No slip condition)<br>2D geometry:<br>- $100 \times 528$ µm or $150 \times 528$ channel geometry for aqueous sample inlet (corresponding to DG250-Y-Mixer or DG300-Y-Mixer, respectively)<br>- $150 \times 1435$ µm channel geometry for oil inlet (DG300-Y-Mixer)<br>- $150 \times 1385$ µm channel geometry for oil inlet (DG250-Y-Mixer)<br>- The aqueous channel joined to the oil channel at a 45° angle.<br>Inlet: Fully Developed Flow (Flow rate in $m^3/s$)<br>Boundary condition = laminar inflow |

| | |
|---|---|
| | Outlet: $p_0 = 0$ |
| Level Set | Surface Domain (Phase initialization):<br>$\frac{\partial \phi}{\partial t} + \boldsymbol{u} \cdot \boldsymbol{\nabla}\phi = \gamma \nabla \cdot \left( \varepsilon \nabla \phi - \phi(1-\phi) \frac{\partial \phi}{\lvert \nabla \phi \rvert} \right)$<br><br>Initial Value and Inlet for the oil phase $\phi$ = 0<br>Initial Value and Inlet for the aqueous phase $\phi$ = 1<br>$\gamma$: 0.0146 m/s<br>$\eta_{oil}$: 6.56 cP<br>$\eta_{aqueous}$: 5.39cP<br>$\rho_{oil}$: 1.8 g/m$^3$<br>$\rho_{aqueous}$: 1 g/m$^3$<br>Surface Tension Coefficient: 14 mN/m<br>Wetted wall: $\theta$ = 2.36 [Rad] for DG 300-Y-Mixer and 2.53 [Rad] for DG250-Y-Mixer. |
| Transport of Diluted Species | Surface Domain<br>$\nabla \cdot (J_i + \boldsymbol{u} c_i) = R_i$ of species $i$<br>$J_i = -D_i \nabla c_i$<br>$c_{substrate}$ = 300 mol/m$^3$<br>$D_{substrate}$ = 6.7*10$^{-10}$ m$^2$/s |
| Nomenclature | $\phi$ = level set function<br>$\boldsymbol{u}$ = Velocity vector of fluid<br>$u$ = fluid velocity [m/s]<br>$p$ = pressure [Pa]<br>$t$ = time [s]<br>$\mu$ = dynamic fluidic viscosity [Pa·s]<br>$c_i$ = Concentration of species $i$ [mol/m$^3$]<br>$c_{sub}$ = Concentration of substrate [mol/m$^3$]<br>$J_i$ = Relative mass flux vector of species $i$<br>$D_i$ = Diffusion coefficient of species $i$<br>$D_{sub}$ = Diffusion coefficient of substrate [m$^2$/s]<br>$\gamma$ = Reinitialization Parameter [m/s]<br>$\eta_{oil}$ = Viscosity of oil [cP]<br>$\eta_{aqueous}$ = Viscosity of aqueous sample (crystal + substrate) [cP]<br>$\rho_{aqueous}$ = Density of aqueous sample [g/m$^3$]<br>$\theta$ = Contact angle [Rad] |
